# Principal component analysis- and tensor decomposition-based unsupervised feature extraction to select more suitable differentially methylated cytosines: Optimization of standard deviation versus state-of-the-art methods

**DOI:** 10.1101/2022.04.02.486807

**Authors:** Y-H. Taguchi, Turki Turki

## Abstract

In contrast to RNA-seq analysis, which has various standard methods, no standard methods for identifying differentially methylated cytosines (DMCs) exist. To identify DMCs, we tested principal component analysis and tensor decomposition-based unsupervised feature extraction with optimized standard deviation, which has been shown to be effective for differentially expressed gene (DEG) identification. The proposed method outperformed certain conventional methods, including those that assume beta-binomial distribution for methylation as the proposed method does not require this, especially when applied to methylation profiles measured using high throughput sequencing. DMCs identified by the proposed method also significantly overlapped with various functional sites, including known differentially methylated regions, enhancers, and DNase I hypersensitive sites. The proposed method was applied to data sets retrieved from The Cancer Genome Atlas to identify DMCs using American Joint Committee on Cancer staging system edition labels. This suggests that the proposed method is a promising standard method for identifying DMCs.

## 1. Introduction

Identification of aberrant methylation (Chen, Lin and Fann, 2016) is important as it plays a critical role in various biological processes. In contrast to the identification of differentially expressed genes (DEGs), effective methods for that of aberrant methylation remain unknown, since it is not directly related to functional units such as genes. There are two approaches to identify aberrant methylation: the identification of differentially methylated cytosines (DMCs) or differentially methylated regions (DMRs). Although these two factors may differ from each other, even if individual cytosines within a region are not differentially methylated and the summation of methylation over the region is differential, it can be considered a DMR. In contrast, even if a region is not a DMR, individual cytosines within the region can be DMCs. This inconsistency provides two ways to identify DMRs. One is a direct method, used for identifying DMRs without identifying individual DMCs. Another is to identify a DMR as an integration of the identified DMCs. Currently, there is no definite consensus on which is the most effective. The identification of DMCs is generally more difficult than that of DMRs due to the huge number of possible methylated CpG sites. For example, there are approximately 28.3 million CpG sites that are predicted to be easily methylated in humans (Luo, Lu and Xie, 2014), which is much larger than the number of human DMRs, which ranges from a few hundred (Skinner, Nilsson, Sadler-Riggleman, Beck, Maamar and McCarrey, 2019) to 20,000 (Shnorhavorian, Schwartz, Stansfeld, Sadler-Riggleman, Beck and Skinner, 2017). As a result, the identification of DMCs is much harder than that of DMRs. To overcome this difficulty, since the microarray era, various computational methods have been proposed to identify DMCs. For example, ChAMP (Tian, Morris, Webster, Yang, Beck, Feber and Teschendorff, 2017) and COHCAP (Warden, Lee, Tompkins, Li, Wang, Riggs, Yu, Jove and Yuan, 2013), developed by illumine, are some of the oldest methods that have been developed to identify DMCs in a specific methylation microarray architecture. For HTS data sets, DMRcate (Peters, Buckley, Chen, Smyth, Goodnow and Clark, 2021), DSS (Park and Wu, 2016), and metilene (Jühling, Kretzmer, Bernhart, Otto, Stadler and Hoffmann, 2015) are some of the most popular methods. However, conventional methods, including these, have common problems that need to be addressed.

1. Conventional methods assume the unrationalized and complicated properties associated with the distribution of methylation.
2. In addition, conventional methods require extended processing times (For example, when we analyzed the GSE77965 data set below, the proposed method required 60 sec for completion whereas ChAMP and COHCAP took 208 sec and 160 sec, respectively. Thus, the conventional methods are two to three times slower).

To address these problems, we applied principal component analysis (PCA)- and tensor decomposition (TD)-based unsupervised feature extraction (FE) with optimized standard deviation (SD) to identify DMCs. This approach was recently successfully applied to identify differentially expressed genes (Taguchi and Turki, 2022). We found that the proposed method can outperform modern methods, especially when applied to HTS data.

## 2. Materials and Methods

### 2.1. Methylation profiles

We used the following four methylation profiles as representatives. We considered profiles previously used for benchmarking or evaluation of previously proposed tools to avoid employing erroneous profiles. In other words, they were previously confirmed to include several DMCs under effective analysis.

#### 2.1.1. GSE77965

This methylation profile was previously used in a benchmark study (BLUEPRINT consortium, 2016) and is available in GEO. From this data, we retrieved six colorectal cancer samples (GSM2062665 to GSM2062670) and six adjacent normal colon tissue samples (GSM2062671 to GSM2062676), and methylation levels were measured using the Illumina HumanMethylation450 BeadChip. Series matrix GSE77965_series_matrix.txt.gz was used.

#### 2.1.2. GSE42308

This methylation profile was used for the evaluation by COHCAP and is available in GEO. It is composed of three HCT116 cell lines and three HCT116 mutants, also measured using the Illumina HumanMethylation450 BeadChip. Series matrix GSE42308_series_matrix.txt.gz was used.

#### 2.1.3. GSE34864

A part of this methylation profile that was originally reported (Smith, Chan, Mikkelsen, Gu, Gnirke, Regev and Meissner, 2012) was used for benchmarking in another study (Feng, Conneely and Wu, 2014) and is available in GEO. We used two oocyte (GSM856495 and GSM856495) and two zygote profiles (GSM856501 and GSM856502) measured by HTS. Four files with names that start with “GSM” and end in ”txt.gz” were used.

#### 2.1.4. EH1072

This methylation profile was included in ExperimentHub (Morgan and Shepherd, 2022) in the Bioconductor and has been used for the evaluation of DMRcate. It has also been analyzed by HTS. It is composed of three Treg cell profiles taken from fat, liver, skin, and lymph nodes and T-cell control taken from lymph nodes (15 samples). For the comparison of conventional methods with the proposed method, since these profiles can only be compared pair-wise, only DMCs between Treg and T-cell control taken from lymph nodes were compared when methods other than the proposed method were employed.

#### 2.1.5. TCGA

Since not all the datasets in TCGA are associated with “ajcc staging system edition”, which was decided to employ in the below. we included the following two TCGA data sets: breast and colon cancer, which are well associated with “ajcc staging system edition”. After selecting files, we downloaded the Manifest file for batch retrieval using gdc-client. Sample and clinical files were also downloaded for the downstream analyses.

To retrieve desired cancer data sets, we used the following parameters in the GDC data portal.

- Data category: dna methylation
- Data Type : Methylation Beta Value
- Platform : illumina human methylation 450

In addition to those defined above, in order to retrieve the specified cancer data set, we used the following parameters for two data sets.

##### Breast

- Primary Site: breast
- project : TCGA-BRCA

this results in 895 files.

##### Colon

- Primary Site: colon
- project : TCGA-COAD

this results in 346 files.

### 2.2. PCA-based unsupervised FE with optimized standard deviation

A full explanation is provided in our previous paper (Taguchi and Turki, 2022) and our recent book (Taguchi, 2020). In brief, the purpose of the method is to select a limited number of variables associated with the classification; in this study, the classifications were clinical, medical, or biological information, e.g., healthy controls and patients. In this specific study, the included variables were the amounts of methylation of individual cytosines within the genome. Since it is unlikely that all cytosine methylation biologically contributes to the difference between classifications, it was expected that a limited number of cytosines would be selected to represent the distinction between classes. This was achieved by finding principal components (PCs) that coincided with the classification and selecting cytosines whose contribution to the corresponding PCs was attributed to individual cytosines. Evaluation of the contribution was performed by relating *P*-values to individual cytosines assuming Gaussian distribution; cytosines with small enough *P*-values were selected as they should have greater contributions than chance alone. See Algorithm S1.

Suppose that *x_ij_* ∈ [0,1]^*N*×*M*^ represents the relative methylation ratio of the *i* th methylation site of the *j* th sample (we assume *N* > *M*). *x_ij_* is normalized as

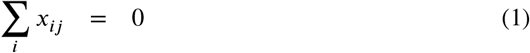

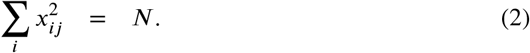

PCA was applied to *x_ij_* so that PC score 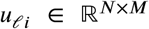 is attributed to the *i*th CpG site and PC loading 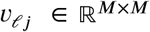 is attributed to the *j*th sample. After identifying PC loading of interest (e.g., distinction between healthy control and patients), we attributed *P*-values to *i*th methylation site assuming that the corresponding PC score, *u_ℓi_*, meets the Gaussian distribution (null hypothesis) as

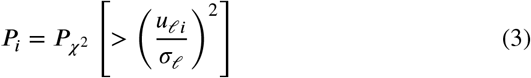

where *P*_*χ*^2^_ [> *x*] is the cumulative *χ*^2^ distribution that the argument is larger than *x* and *σ_ℓ_* is the standard deviation optimized as much as possible to obey the null hypothesis (see below). We assumed a Gaussian distribution for *u_ℓ_i__*. for several reasons: Firstly, it worked reasonably well in previous studies (Taguchi, 2020). Secondly, a Gaussian distribution is expected to occur when random numbers are added because of the central limit theorem. Thus, we assumed the null hypothesis that *u_ℓ_i__* obeys the Gaussian distribution, and it performs well, as demonstrated below. When the histogram of 1 – *P_i_* obeys the null hypothesis, it should be flat, excluding a sharp peak located near 1 – *P_i_* ~ 1, which represents is to be selected. The optimization of *σ_ℓ_* is performed with minimization of the standard deviation of 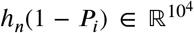 for microarray, 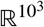 for HTS, *n* < *n*_0_ where *h_n_* represents *n*th histogram and *n*_0_ is the largest *n* that satisfies the adjusted *P_i_*(∈ *h_n_*) that is larger than *P*_0_, a threshold adjusted *P*-value.

The optimization of *σ_ℓ_* was as follows: To minimize the SD of binned *h_n_* = *h*(1 – *P_i_*) *σ_h_*,

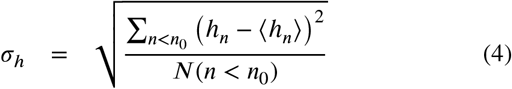

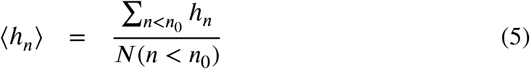

with respect to *σ_ℓ_*, where *N*(*n* < *n*_0_) is the number of *n*s associated with *n* < *n*_0_, i.e., they are not recognized as outliers but are recognized as a part of the Gaussian distribution; thus, we selected the *σ_ℓ_* in eq. (3) that resulted in the smallest *σ_h_*. *P*-values are corrected using the BH criterion (Taguchi, 2020) and *i*th associated with adjusted *P*-values less than *P*_0_, which is typically 0.01, are selected as DMCs.

### 2.3. TD-based unsupervised FE with optimized standard deviation

The purpose of the method is similar to the previous section, i.e., selecting variables that contribute to the classification. However, the computational method is different as PCA is replaced with TD. See also Algorithm S2. A full explanation is, again, provided in our previous paper (Taguchi and Turki, 2022) and our recent book (Taguchi, 2020). As TD was only applied to EH1072 here, we assumed that the tensor represented the EH1072 dataset.

Suppose 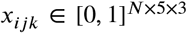 represents methylation of *i* cytosine in *j*th cells (*j* = 1: Treg cells from fat, *j* = 2: Treg cells from liver, *j* = 3: Treg cells from skin, *j* = 4:Treg cells from lymph nodes, *j* = 5:T-cells from lymph nodes) of the *k*th biological replicate. *x_ijk_* is normalized as

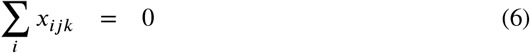

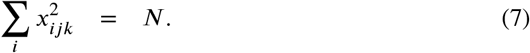

We applied higher order singular value decomposition, HOSVD (Taguchi, 2020), to *x_ijk_* and obtained TD as

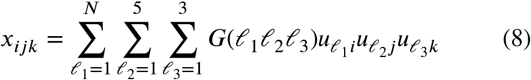

where 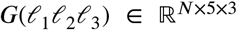 is core tensor, and 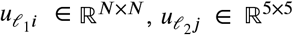 and 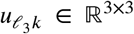 are singular value matrices and orthogonal matrices, respectively. To select the *u*_*ℓ*_1_*i*_ used to detect DMCs, we selected *u*_*ℓ*_2_*j*_ and *u*_*ℓ*_3_*k*_, which had the desired properties of dependence upon cells and independence from replicates. After identifying the *u_ℓ_2_j_* and *u*_*ℓ*_3_*k*_ needed, we checked 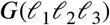 to see which *ℓ_1_* shares *G*s that have the largest absolute values, with the selected *ℓ*_2_ and *ℓ*_3_. Finally, *P*-values were attributed to *i* with the equation

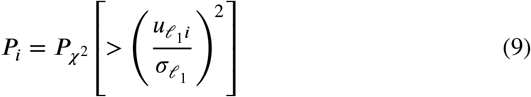

after optimizing *σ*_*ℓ*_1__, as described in the previous subsection. *P*-values were corrected using the BH criterion and *i*th was associated with adjusted *P*-values that were less than *P*_0_, which was typically 0.01, were selected as DMCs.

### 2.4. Overlap between DMCs and various functional regions for microarray

The microarray used in this study was the Illumina HumanMethylation450, the annotation of which is available as GPL16304 from GEO. Among the various functional regions annotated, we considered known DMRs, enhancers, and DHS. Fisher’s exact test was applied to see if DMCs significantly overlapped with DMRs, enhancers, or DHS.

### 2.5. Identification of coincidence between selected DMC and DHS for HTS

To compare selected DMCs with DHSs, we downloaded http://mirror.vcu.edu/vcu/encode/DNase-seq_Peaks/wgEn-codeDukeDnaseFibrobl.fdr01peaks.hg19.bb, which is the fibroblast DHS in the hg19 genome, to which short reads were mapped from HTS data that were investigated in this study (i.e., GSE34864 and EH2072). *t* test was applied to *P*-values obtained by the proposed method to see if there were distinctions between DHSs and non-DHSs in individual chromosomes.

### 2.6. eDMR

eDMR was performed with the options ACF=F, DMR. methdiff=1, and DMC.methdiff=1, using the normalized third PC score 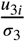 (for GSE34864) and the normalized second PC score 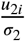 (for EH1072) as meth.diff where *σ_ℓ_* is the optimized SD. Then, filter.dmr was executed with the options DMR. qvalue =0.01 and mean.meth.diff=1. Other options were set as the default.

### 2.7. metilene

Metilene was performed with the default options.

### 2.8. Binomial test for the overlap between two genomic regions

We considered two genomic regions, *g*_1_ and *g*_2_, whose associated sequence lengths are *ℓ*_*g*_1__ and *ℓ*_*g*_2__, respectively, within the whole genome *g*, associated with the sequence length *ℓ*_g_. In *g*_1_, the region with length 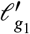 at least partially overlapped with *g*_2_. Similarly, the region with length 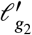 in *g*_2_ at least partially overlapped with *g*_1_. We evaluated the significance of this overlap using the cumulative binomial distribution *P_b_*(> *x, N, p*), which represents a cumulative probability where the argument is larger than *x* (which is selected within *N*) and with an individual probability of the selection *p* for 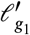 and 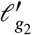 as

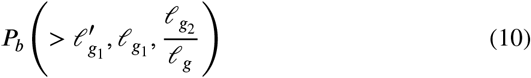

and

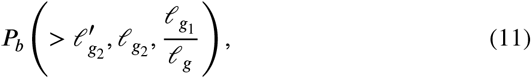

respectively.

### 2.9. Enrichment analysis

Enrichment analysis was performed using Enrichr (Kuleshov, Jones, Rouillard, Fernandez, Duan, Wang, Koplev, Jenkins, Jagodnik, Lachmann, McDermott, Monteiro, Gundersen and Ma’ayan, 2016). Gene symbols associated with selected probes were identified in GPL13534, which was downloaded from GEO (GEO ID: GPL13534). The numbers of probes considered were also restricted when COHCAP was applied, since COHCAP identified too many probes as DMCs. *P*-values computed by COHCAP were used to rank identified probes. Additionally, only top ranked probes were considered so as to have as many gene symbols as those identified by the proposed method. Then, obtained gene symbols were uploaded to Enrichr.

## 3. Results

### 3.1. Microarray

First, we applied the proposed method to methylation profiles measured by microarray architecture. PCA was applied to *x_ij_* of GSE77965. Then, we found that *v*_2*j*_ was associated with the distinction between two classes (Fig. S1 (a), *P* = 5.68×10^−3^ computed by *t* test). Then, we attempted to optimize *σ*_2_ so that *u*_2*i*_ met the Gaussian distribution as much as possible. We found that *σ*_2_ = 0.1506726 can result in the smallest standard deviation of *h_n_* (Fig. S2 (a)). Indeed, the histogram of 1 – *P_i_* was close to the ideal case when *σ*_2_ the optimized value is used (Fig. S2 (a)). *P_i_*s were corrected using the BH criterion. We found that 87,665 methylation sites were associated with adjusted *P*-values less than 0.01 and were regarded as DMCs. To evaluate whether the selected DMCs were reasonable, we compared them with the information attached to the microarray (i.e., annotations attached to microarray probes). The high coincidence with functional sites (Table S1(a)) validated our selection of DMCs.

The same process was applied to GSE42308. We, again, found that the second PC was associated with two classes (Fig. S1(b), *P* = 9.21 × 10^−7^ computed by *t* test). We also attempted to optimize *σ*_2_ so that *u*_2*i*_ met a Gaussian distribution as much as possible. We found that *σ*_2_ = 0.07036153 resulted in the smallest standard deviation of *h_n_* (Fig. S2 (b)). In actuality, the histogram of 1 – *P_i_* was close to the idealized case when *σ*_2_ is the optimized value (Fig. S2(b)). *P_i_*s were corrected using the BH criterion. We found that 97,091 methylation sites were associated with adjusted *P*-values less than 0.01 and could be regarded as DMCs. To evaluate if these selected DMCs were reasonable, we compared them with information attached to the microarray (i.e., annotations attached to microarray probes). Their high coincidence with the functional sites (Table S1 (b)) further proved our selection of DMCs to be reliable. However, the number of false positives obtained was not negligible; thus, further improvement is evidently required.

### 3.2. HTS

Next, we attempted to validate the effectiveness of the proposed method in not only microarrays, but also HTS. To this end, we applied the proposed method to GSE34864. Unfortunately, owing to the small number of samples (4), no PC loading attributed to samples *j* showed significant distinction between the two classes. Nevertheless, since DMCs appeared to be identifiable using the proposed method, we employed *ℓ* = 3 as it was most associated with a distinction between two classes (Fig. S1(c)). We also tried to optimize *σ*_3_ so that *u*_3*i*_ met the Gaussian distribution as much as possible. We found that *σ*_3_ = 0.3499989 resulted in the smallest standard deviation of *h_n_* (Fig. S2 (c)). In actuality, the histogram of 1 – *P_i_* was close to the idealized case when *σ*_3_ takes the optimized value (Fig. S2(c)). In contrast to the microarray results, we lacked annotations for methylation sites. Therefore, we checked the coincidence between DHSs and identified DMCs. We applied a *t* test to assess the null hypothesis that the distribution of *P_i_* was identical between DHS and non-DHS sites in individual chromosomes (Figure S3(a)). Thereafter, we found *P* = 0 for all chromosomes within numerical accuracy. Thus, it was obvious that the proposed method could identify DMCs correctly. However, the number of false positives obtained was not negligible. Further improvement is, hence, required.

Then, we applied the proposed method to yet another HTS dataset, EH1072. After applying PCA, we found that *v*_2*j*_ was associated with distinction between five cell classes (Fig. S1(d), *P* = 2.22 × 10^−3^ computed by categorical regression). We also tried to optimize *σ*_2_ so that *u*_2*i*_ met the Gaussian distribution as much as possible. We found that *σ*_2_ = 0.2846131 could result in the smallest standard deviation of *h_n_* (Fig. S2(d)). Indeed, the histogram of 1 – *P_i_* was close to the ideal case when *σ_3_* took the optimized value (Fig. S2(d)). In contrast to the cases for the microarray analysis, we lacked annotations for methylation sites. Therefore, we checked the coincidence between DHSs and identified DMCs. We applied a *t* test to assess the null hypothesis that the distribution of *P_i_* was identical between the DHS and non-DHS sites in individual chromosomes. We found *P* = 0 for all chromosomes with numerical accuracy (Figure S3(b)). Thus, the proposed method could identify DMCs accurately.

Finally, to test the effectiveness of TD, we applied ”TD-based unsupervised FE with optimized SD” to EH1072. We also applied HOSVD to *x_ijk_*; Fig. S1(e) shows the first and second *u*_*ℓ*_2_*j*_ attributed to *j*th cells and the first *u*_*ℓ*_3_*k*_ attributed to the *k*th biological replicate. Since *u*_2*j*_ represented the dependence upon cells similar to that seen in Fig. S1 (d) and *u*_1*k*_ represented the independence of biological replicates, we needed to identify which *u*_*ℓ*_1_*i*_ attributed to the *i*th cytosine site was associated with *u*_2*j*_ and *u*_1*k*_ (for other *u*_*ℓ*_2_*j*_ and *u*_*ℓ*_3_*k*_s, see Figs. S4 and S5, which clearly indicated that no other singular value vectors were associated with the desired properties).

Table S2 shows *G*(*ℓ*_1_, 2, 1) which takes the largest absolute value for *ℓ*_1_ = 2. Thus, although *u*_2*i*_ should be used to select DMCs, it is equal to the second PC score attributed to *i* since HOSVD is mathematically equivalent to SVD applied to unfolded tenor (Taguchi, 2020). Therefore, TD will select the same set of DMCs as PCA. This implies that TD works equally as well in DMC identification.

### 3.3. Comparisons with other methods

To evaluate the effectiveness of the proposed method, we compared its performance with that of other methods. Although there is a comprehensive review (Chen et al., 2016), we selected COHAMP for microarray architecture and DSS for HTS architecture as representatives from these reviews. As newer ones were not included in this review, we additionally selected ChAMP for microarray architecture and DMRcate and metilene for HTS architecture.

Firstly, we considered the application of the proposed model for microarray architecture. The first method to be compared with the proposed method was COHCAP. First, we applied COHCAP to GSE77965. Table S1 (c) shows the coincidence between the selected and functional regions. When compared with the performance of the proposed method (Table S1(a)), the performance was generally worse. Next, we applied COHCAP to GSE42308. Table S1(d) shows the coincidence between the selected and functional regions. When compared with the performance of the proposed method (Table S1(b)) the performance was slightly better; however, the number of DMRs was much lower than that obtained using the proposed method. In conclusion, the comparisons of GSE77965 and GSE42308 suggest that the overall performance of the proposed method was better than that of COHCAP. To explore whether the superiority of the proposed method is accidental, we also compared it to another method, ChAMP. Table S1(e) shows the coincidence between the selected and functional regions. When compared with the performance of the proposed method (Table S1(a)), although the performance was comparable, the number of regions selected by the proposed method was roughly ten times greater than that selected by ChAMP. Thus, the proposed method can identify more DMRs containing more identified DMCs while retaining a significant overlap with functional regions. Next we applied ChAMP to GSE42308. Table S1(f) shows the coincidence between the selected and functional regions. When compared with the performance of the proposed method (Table S1(b)), the performance and the number of DMRs were comparable. The comparisons using GSE77965 and GSE42308 suggest that the overall performance of the proposed method was better than that of COHCAP, as well as ChAMP. Thus, the superiority of the proposed method, when applied to microarray architecture, over the conventional method was unlikely due to chance. Next, to observe the performance of the proposed model with HTS data, we first tested DSS. For GSE34864, we found that *P* = 0 for the *t* test denies the probability that the distribution of *P*-values attributed to DHS and non-DHS sites was identical (Figure S3(c)). For EH1072, since DSS required more than a week for processing, even on using 10 CPU cores, we terminated the computation. We next applied DMRcate to GSE34864 and EH1072. For GSE34864, DMRcate failed to identify any DMCs. For EH1072, we found that *P = 0* for the *t* test denies the probability that the distribution of *P*-values attributed to DHS and non-DHS sites was identical (Figure S3(d)). We also employed metilene, which failed to identify DMCs for either GSE34864 or EH1072. Thus, the overall performance of the proposed method was better than DSS, DMRcate, and metilene. The use of epigenome-wide association studies (EWAS) (Flanagan, 2014) was outside the scope of this model since EWAS requires numerous samples, while the proposed method was effective when the number of samples was less than ten.

Although the effectiveness of the proposed method is questionable since it could not identify DMRs for HTS architecture, we could identify DMRs by integrating identified DMCs. To show this ability, we used eDMR (Li, Garrett-Bakelman, Akalin, Zumbo, Levine, To, Lewis, Brown, D’Andrea, Melnick and Mason, 2013) to identify DMRs by integrating identified DMCs. We found that eDMR identified 6,818 and 12,662 DMRs for GSE34864 and EH1072, respectively. Thus, the proposed method may also have the ability to identify DMRs if a suitable tool is employed. In contrast, 27 and 12,478 DMRs were identified by metilene for GSE34864 and EH1072, respectively, whilst it failed to identify DMCs for these profiles. Thus, eDMR combined with the proposed method identified more DMRs than metilene.

To confirm the significant overlap of genomic regions selected by eDMR with DHS, we used a binomial distribution for the evaluation (see Methods); we could not employ Fisher’s exact test for the HTS, and since eDMR did not attribute *P*-values to DMRs, we could not employ a *t* test.

Table S3 shows *P*-values that are 0 within numerical accuracy. Thus, DMRs selected by eDMR significantly overlapped with DHS.

### 3.4. Application to TCGA

Above, we have successfully demonstrated the superiority of the proposed method over the conventional methods. In this subsection, we discuss the application of studies associated with biological evaluation using enrichment analyses to strengthen the efficiency of the proposed method. We employed TCGA (see Acknowledgement), which contains versatile cancer data sets. One possible limitation of TCGA is that its DNA methylation data is rarely studied independently; it is usually analyzed in combination with additional data, e.g., gene expression data. Since the purpose of the present study was to demonstrate the effectiveness of the proposed method when applied to *only* DNA methylation alone, integrated analysis with other data is out of its scope. However, to overcome this challenge and to increase the novelty of this method, we have made the following additions. First of all, we used “ajcc staging system edition” (Edge and Compton, 2010) as clinical information. The American Joint Committee on Cancer (AJCC) releases the staging labels of cancer development. Thus, the staging system is a good indicator of the grades of individual samples.

Among the methods compared with the proposed methods, ChAMP and COHCAP can be applied to TCGA, since TCGA employs microarray architecture. Between ChAMP and COHCAP, only COHCAP be used for “ajcc staging system edition”, since it has multiple levels, whereas ChAMP is not suitable for analysis of multiple classes, only pairwise comparisons.

For our results, for both breast and colon data sets, the second PC attributed to samples, *v*_2*j*_, was significantly associated with “ajcc staging system edition”. Then, the PC attributed to sites, *u*_2*i*_, was employed and SD was optimized. As a result, we identified 9,544 and 11,447 probes that were associated with distinctions between multiple classes in “ajcc staging system edition”. COHCAP was also applied to the two data sets, and 333,949 and 21,042 probes were selected that were associated with distinctions between multiple classes in “ajcc staging system edition” (when multiple classed were considered, COHCAP utilized ANOVA to select sites). Since the number of probes selected by COHCAP was very large, it is unlikely that all of them were differentially methylated sites. Thus, we restricted the number of probes selected by COHCAP to obtain as many associated gene symbols as those identified by the proposed method (see below). The numbers of gene symbols associated with the probes selected by the proposed method were 4,041 and 2,815 for breast and colon data sets, respectively. Those selected by COHCAP were restricted to 4,326 and 2,974 for breast and colon data sets, respectively, by reducing the numbers of probes to the 6,000 and 5,000 top ranked probes associated with smaller *P*-values for breast and colon data sets, respectively. We have uploaded the four lists of gene symbols associated with identified methylation sites to Enricher (Kuleshov et al., 2016). The following are some examples of categories associated with enriched terms associated with the identified gene symbols. Figure 1 shows the results for the “Cancer Cell Line Encyclopedia” category of Enricher. The proposed method identified significant cell lines within the top 10 ranked cell lines whereas COHCAP failed (no cell lines associated with significant adjusted *P*-values less thna 0.05). This is only one example but there could be more cases where the proposed method is successful and COHCAP fails (i.e., the proposed method could identify the top 10 significant cell lines but COHCAP could not; Figs. S6 for “ARCHS4 Cell-lines”, S7 for “KEGG 2021 HUMAN”, S8 for “GO Biological process 2021”, S9 for “Jensen Diseases”, S10 for “GO Molecular function 2021”, S11 for “GO Cellular Components 2021”), which are summarized in Table 1. Thus, as least for some cases, the proposed method can outperform COHCAP from a biological point of view.

**Figure 1:**
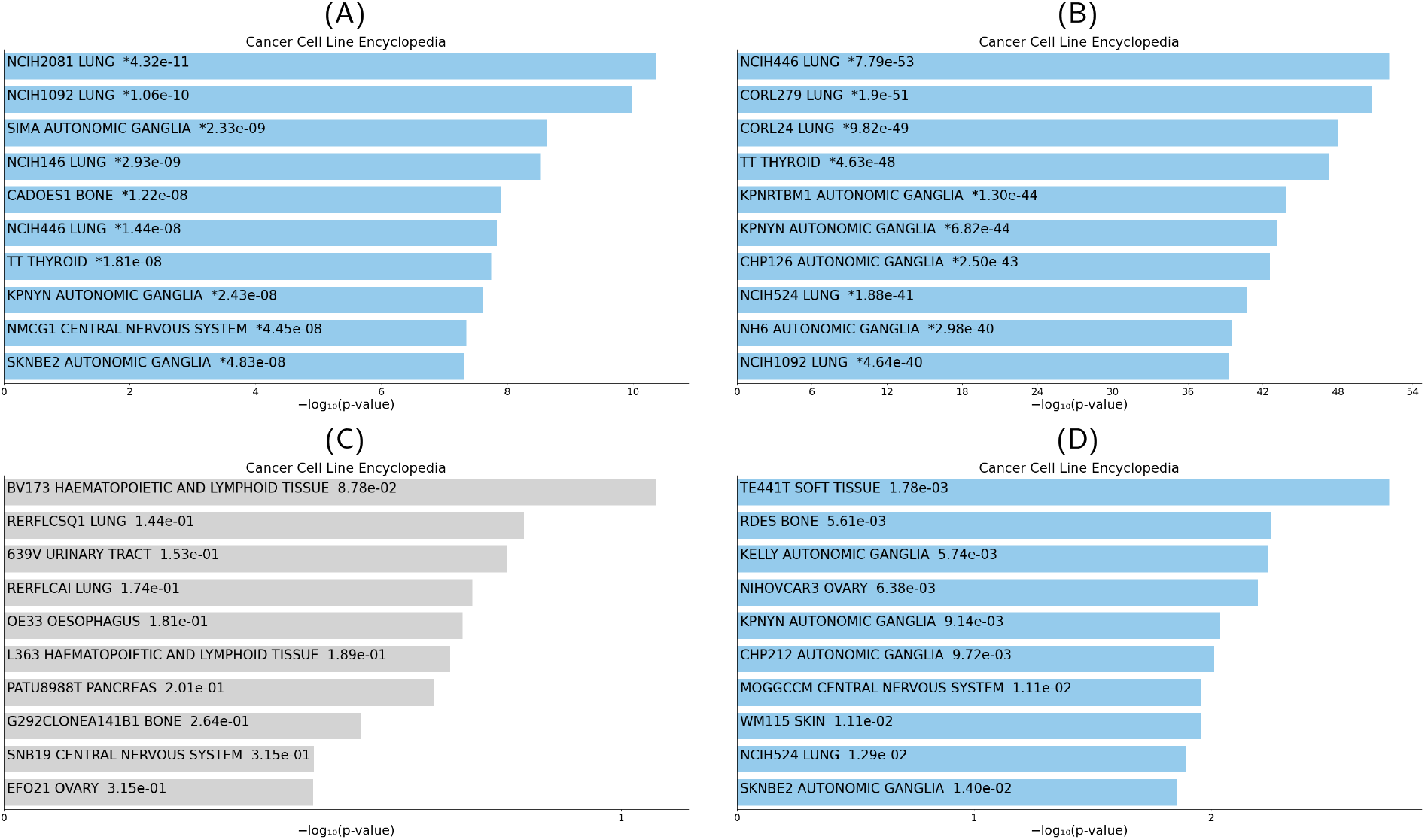
Top 10 ranked cell lines in the “Cancer Cell Line Encyclopedia” category of Enricher for genes selected by the proposed method for (A) breast and (B) colon data sets and those selected by COHCAP for (C) breast and (D) colon data sets, visualized by Appyters (Clarke et al., 2021). Blue colored: raw *P* < 0.05 and asterisked: adjusted *P* < 0.05, respectively.

**Table 1.** Number of significant (i.e., adjusted *P*-values are less than 0.05) variables (e.g., cell lines, pathways, and diseases e.t.c.) among the top 10 ranked factors in individual categories for the proposed method and COHCAP

| Category | Proposed method |  | COHCAP |  | Figs. |
| --- | --- | --- | --- | --- | --- |
|  | Breast | Colon | Breast | Colon |  |
| Cancer Cell Line Encyclopedia | 10 | 10 | 0 | 0 | 1 |
| ARCHS4 Cell-lines | 10 | 10 | 0 | 0 | S6 |
| KEGG 2021 HUMAN | 10 | 10 | 2 | 0 | S7 |
| GO Biological process 2021 | 10 | 10 | 1 | 9 | S8 |
| Jensen Diseases | 10 | 10 | 3 | 7 | S9 |
| GO Molecular function 2021 | 10 | 10 | 6 | 7 | S10 |
| GO Cellular components 2021 | 9 | 10 | 6 | 0 | S11 |

## 4. Discussion

Although TD- and PCA-based unsupervised FE without optimized SD were developed many years ago and have been successfully applied to a wide range of studies (Taguchi, 2020), the novelty of this study is that we have advanced the method by optimizing the SD so that the histogram of 1 – *P_i_* obeys the null hypothesis as much as possible. This advancement is significant since TD- and PCA-based unsupervised FE, without optimized SD, could only identify a very small number of DMCs. Furthermore, it is unlikely that there are no false negatives and the histogram of 1 – *P_i_* is far from optimal. The optimization of SDs results in a more ideal histogram of 1 – *P_i_* and a sufficient number of DMCs, which competes with state-of-the-art methods, as shown above. In addition, it is rare that a tool can be applicable to both gene expression and DNA methylation measurements by both microarray analysis and HTS without any specific modifications. For example, although Rackham, Dellaportas, Petretto and Bottolo (2015) published a benchmark of DNA methylation tools, no tools specific to gene expression were included. This implies that tools specific to gene expression are not expected to work well for DNA methylation as well. Nevertheless, the proposed method (previously tested for gene expression (Taguchi and Turki, 2022)) was effective for DNA methylation without any specific modifications. Hence, our reporting of this remarkable finding. Thus, although this advancement may be small, the improvement in outcomes will be large.

As can be seen in Figs. S2 (a), (b), (c) and (d), the SD of *h_n_* becomes zero when the optimized SD is equal to zero. This is inefficient since all *i*s have *P* = 0 and are selected as DMCs. This results in an empty {*n* < *n*_0_} since no is have a larger *P_i_* than the threshold value *P*_0_, no matter how small *P*_0_ is. Thus, we need to find the smallest SD to have the smallest SD of *h_n_* with an excluding SD of zero.

We employed a *t* test for HTS instead of a Fisher’s exact test as, for microarray architecture, the selection was performed among probes with annotations. Thus, we could compare two classifications such as functional probes and non-functional probes versus selected probes and nonselected probes. Nevertheless, for HTS, since there was only a list of DHSs and DMCs, there were no common background sets from which we could define non-selected and non-DHS sites. Thus, we could not perform a Fisher’s exact test, and opted to apply a *t* test to *P*-values between DHS and non-DHS as this was possible even without a common background set.

In conclusion, in this paper, we applied the proposed method “PCA- and TD-based unsupervised FE with optimized SD” to identify DMCs. Identified DMCs were significantly overlapped with various functional sites. This technique also outperformed some modern methods, especially when applied to HTS architecture. We also identified DMRs by integrating DMCs. The proposed method does not have to assume non-trivial distribution such as the beta-binomial distribution for methylation and, instead, simply assumes that the PC score obeys a Gaussian distribution, which is rationalized in the framework for probabilistic PCA (Tipping and Bishop, 1999); therefore, the proposed method is more adequate than other current methods that must assume betabinomial distribution.

## Supporting information

Supplementary Document(Algorithm)

## CRediT authorship contribution statement

**Y-H. Taguchi:** Planning the research, Analyses, Valuation of the results, discussions, and outcomes, Wring the manuscript. **Turki Turki:** Valuation of the results, discussions, and outcomes, Wring the manuscript.

## Institutional Review Board Statement

Not applicable.

## Informed Consent Statement

Not applicable.

## Data availability

All data used in this study are cited in the manuscript and are available by public access. Sample R source code is available at https://github.com/tagtag/PCAUFEOPSD/ (Accessed 5th January 2023).

## Acknowledgement

The results shown here are in part based upon data generated by the TCGA Research Network: https://www.cancer.gov/tcga.

