## Supplementary Tables for "Principal component analysis- and tensor decomposition-based unsupervised feature extraction to select more suitable differentially methylated cytosines: Optimization of standard deviation versus state-of-the-art methods"

Table S1: Fisher's exact tests between identified DMCs and various annotations for (a) GSE77965, (b) GSE42308, COHCAP applied to (c) GSE77965, (d) GSE42308, ChAMP applied to (e) GSE77965, (f) GSE42308. DMR: Differentially methylated region, DHS: DNase I hypersensitive site

(a) GSE77965

| | adjusted $P_i$ | |
|---|---|---|
| DMR | $> 0.01$ | $< 0.01$ |
| NO | 378862 | 69378 |
| YES | 19050 | 18287 |
| odds ratio | 5.24 | |
| $P$-value | 0 | |

| | adjusted $P_i$ | |
|---|---|---|
| Enhancer | $> 0.01$ | $< 0.01$ |
| NO | 321561 | 61457 |
| YES | 76351 | 26208 |
| odds ratio | 1.80 | |
| $P$-value | 0 | |

| | adjusted $P_i$ | |
|---|---|---|
| DHS | $> 0.01$ | $< 0.01$ |
| NO | 352206 | 73455 |
| YES | 45706 | 14210 |
| Odds ratio | 1.49 | |
| $P$-value | $8.53 \times 10^{-305}$ | |

(b) GSE42308

| | adjusted $P_i$ | |
|---|---|---|
| DMR | $> 0.01$ | $< 0.01$ |
| NO | 361067 | 87173 |
| YES | 27419 | 9918 |
| odds ratio | 1.50 | |
| $P$-value | $4.77 \times 10^{-224}$ | |

| | adjusted $P_i$ | |
|---|---|---|
| Enhancer | $> 0.01$ | $< 0.01$ |
| NO | 313295 | 69723 |
| YES | 75191 | 27368 |
| odds ratio | 1.64 | |
| $P$-value | 0 | |

| | adjusted $P_i$ | |
|---|---|---|
| DHS | $> 0.01$ | $< 0.01$ |
| NO | 343294 | 82367 |
| YES | 45192 | 14724 |
| odds ratio | 1.36 | |
| $P$-value | $2.37 \times 10^{-188}$ | |

(c) GSE77965 (COHCAP)

| | adjusted $P_i$ | |
|---|---|---|
| DMR | $> 0.01$ | $< 0.01$ |
| NO | 447353 | 887 |
| YES | 37233 | 104 |
| odds ratio | 1.41 | |
| $P$-value | $1.51 \times 10^{-3}$ | |

| | adjusted $P_i$ | |
|---|---|---|
| Enhancer | $> 0.01$ | $< 0.01$ |
| NO | 382429 | 589 |
| YES | 102157 | 402 |
| odds ratio | 2.55 | |
| $P$-value | $1.62 \times 10^{-43}$ | |

| | adjusted $P_i$ | |
|---|---|---|
| DHS | $> 0.01$ | $< 0.01$ |
| NO | 424801 | 860 |
| YES | 59785 | 131 |
| odds ratio | 1.08 | |
| $P$-value | $4.11 \times 10^{-1}$ | |

(d) GSE42308 (COHCAP)

| | adjusted $P_i$ | |
|---|---|---|
| DMR | $> 0.01$ | $< 0.01$ |
| NO | 434162 | 14078 |
| YES | 35215 | 2122 |
| odds ratio | 1.86 | |
| $P$-value | $5.86 \times 10^{-129}$ | |

| | adjusted $P_i$ | |
|---|---|---|
| Enhancer | $> 0.01$ | $< 0.01$ |
| NO | 371646 | 11372 |
| YES | 97731 | 4828 |
| odds ratio | 1.61 | |
| $P$-value | $8.38 \times 10^{-154}$ | |

| | adjusted $P_i$ | |
|---|---|---|
| DHS | $> 0.01$ | $< 0.01$ |
| NO | 412252 | 13409 |
| YES | 57125 | 2791 |
| odds ratio | 1.50 | |
| $P$-value | $3.61 \times 10^{-75}$ | |

(e) GSE77965(ChAMP)

| | adjusted $P_i$ | |
|---|---|---|
| DMR | $> 0.01$ | $< 0.01$ |
| NO | 442353 | 5887 |
| YES | 35069 | 2268 |
| odds ratio | 4.86 | |
| $P$-value | 0 | |

| | adjusted $P_i$ | |
|---|---|---|
| Enhancer | $> 0.01$ | $< 0.01$ |
| NO | 377718 | 5300 |
| YES | 99704 | 2855 |
| odds ratio | 2.04 | |
| $P$-value | $1.18 \times 10^{-186}$ | |

| | adjusted $P_i$ | |
|---|---|---|
| DHS | $> 0.01$ | $< 0.01$ |
| NO | 419579 | 6082 |
| YES | 57843 | 2073 |
| odds ratio | 2.47 | |
| $P$-value | $5.50 \times 10^{-231}$ | |

(f) GSE42308(ChAMP)

| | adjusted $P_i$ | |
|---|---|---|
| DMR | $> 0.01$ | $< 0.01$ |
| NO | 374324 | 73916 |
| YES | 28338 | 8999 |
| odds ratio | 1.61 | |
| $P$-value | $1.86 \times 10^{-283}$ | |

| | adjusted $P_i$ | |
|---|---|---|
| Enhancer | $> 0.01$ | $< 0.01$ |
| NO | 324955 | 58063 |
| YES | 77707 | 24852 |
| odds ratio | 1.79 | |
| $P$-value | 0 | |

| | adjusted $P_i$ | |
|---|---|---|
| DHS | $> 0.01$ | $< 0.01$ |
| NO | 356284 | 69377 |
| YES | 46378 | 13538 |
| odds ratio | 1.50 | |
| $P$-value | $1.01 \times 10^{-301}$ | |

Table S2: $G(\ell_1, 2, 1)$ in eq. (8 ) with $\ell_2 = 2$ and $\ell_3 = 1$.

| $\ell_1$ | $G(\ell_1, 2, 1)$ | $\ell_1$ | $G(\ell_1, 2, 1)$ |
|---|---|---|---|
| 1 | -10 | 6 | 44 |
| 2 | 2949 | 7 | 7 |
| 3 | 6 | 8 | 31 |
| 4 | 5 | 9 | -22 |
| 5 | -18 | 10 | 12 |

Table S3: Binomial test for the coincidence between DHS and DMRs, identified by eDMR.

| | | GSE34864 | EH1072 |
|---|---|---|---|
| | Human genome ($\ell_g$) | 3,117,275,501 | |
| | DHS ($\ell_{g_1}$) | 61,495,656 | |
| Length of | Overlap ($\ell'_{g_1}$) | 86,523 | 53,907 |
| | DMR ($\ell_{g_2}$) | 3,246,111 | 742,103 |
| | Overlap ($\ell'_{g_2}$) | 418,206 | 25,541 |
| | $P_b\left(> \ell'_{g_1}, \ell_{g_1}, \frac{\ell_{g2}}{\ell_g}\right)$ | 0 | 0 |
| | $P_b\left(> \ell'_{g_2}, \ell_{g_2}, \frac{\ell_{g_1}}{\ell_g}\right)$ | 0 | 0 |
