## Supplementary Figures for "Principal component analysis- and tensor decomposition-based unsupervised feature extraction to select more suitable differentially methylated cytosines: Optimization of standard deviation versus state-of-the-art methods"

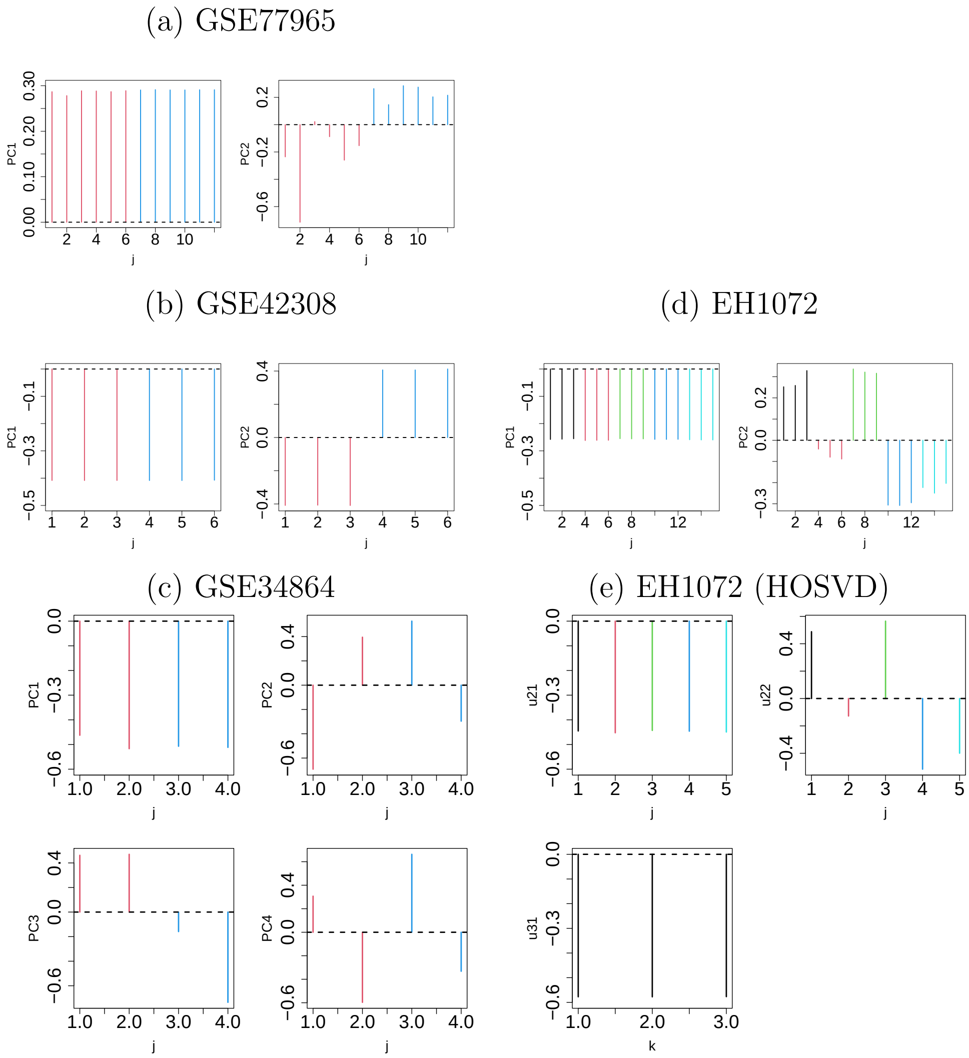


Figure S1: The first and second PC scores ($v_{1j}$ and $v_{2j}$) attributed to the $j$th sample for (a) GSE77965, Red: tumors, blue: normal tissues, (b) GSE42308, Red: HCT116 cell lines, blue: HCT116 mutants, (c) GSE34864, Red: oocytes, blue: zygote, (d) EH1072, Black: Treg from fat, red: Treg from liver, green: Treg from skin, blue: Treg from lymph node, cyan: Treg control from lymph node. (e) Upper: The first and second $u_{\ell_2 j}$ attributed to $j$th cells. Black: Treg from fat, red: Treg from liver, green: Treg from skin, blue: Treg from lymph nodes, cyan: Treg control from lymph node. Lower: The first $u_{\ell_3 k}$ attributed to biological replicates.

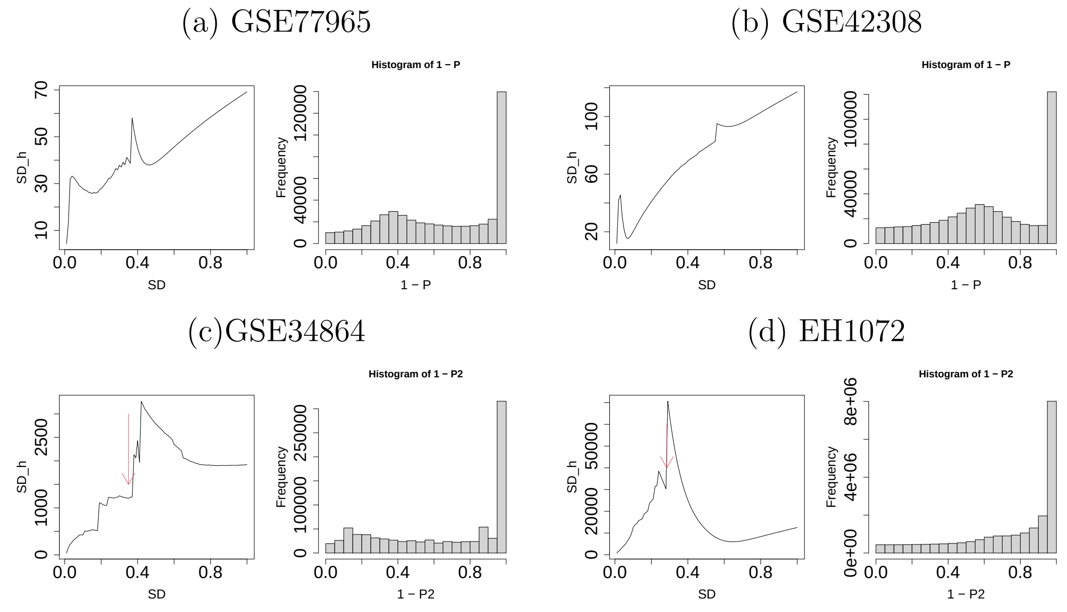


Figure S2: Left: Dependence of standard deviation of $h_n$ on $\sigma_2$. Right: Histogram of $1 - P_i$ for the optimized $\sigma_2$. (a) GSE77965, (b) GSE42308, (c) GSE34864, (d) EH1072.

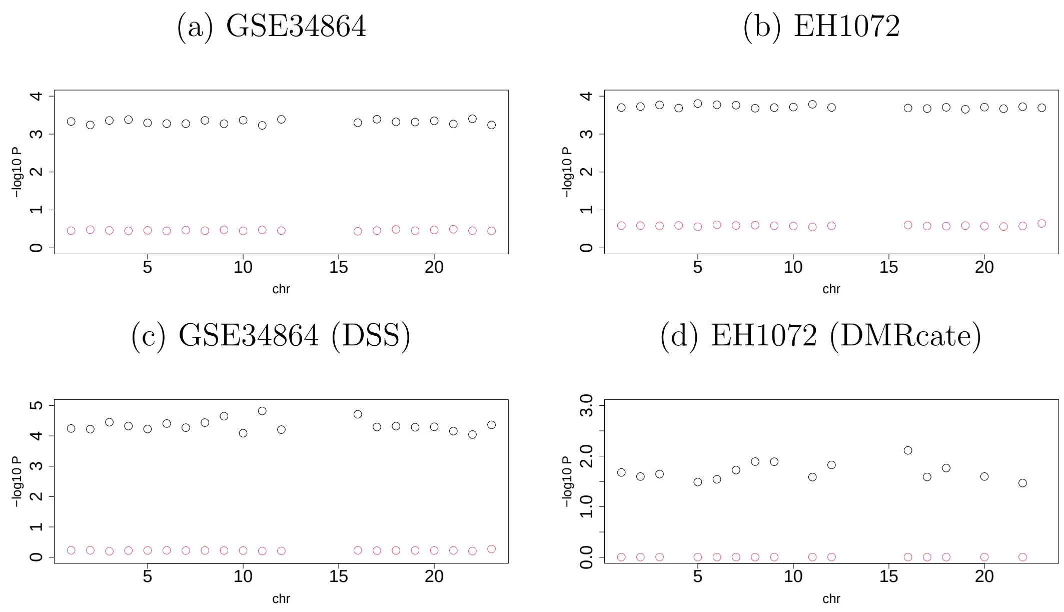


Figure S3: Negative logarithm of mean *P*-values between DHS (black circles) and non-DHS sites (red circles) within each chromosome (horizontal axis) when PCA-based unsupervised FE with optimized SD was applied to (a)GSE34864, (b) EH1072, (c) DSS applied to GSE34864, (d) DMRcate applied to EH1072. *P*-values computed using $t$ test between these two were zero. Some chromosomes did not have an associated DHS.

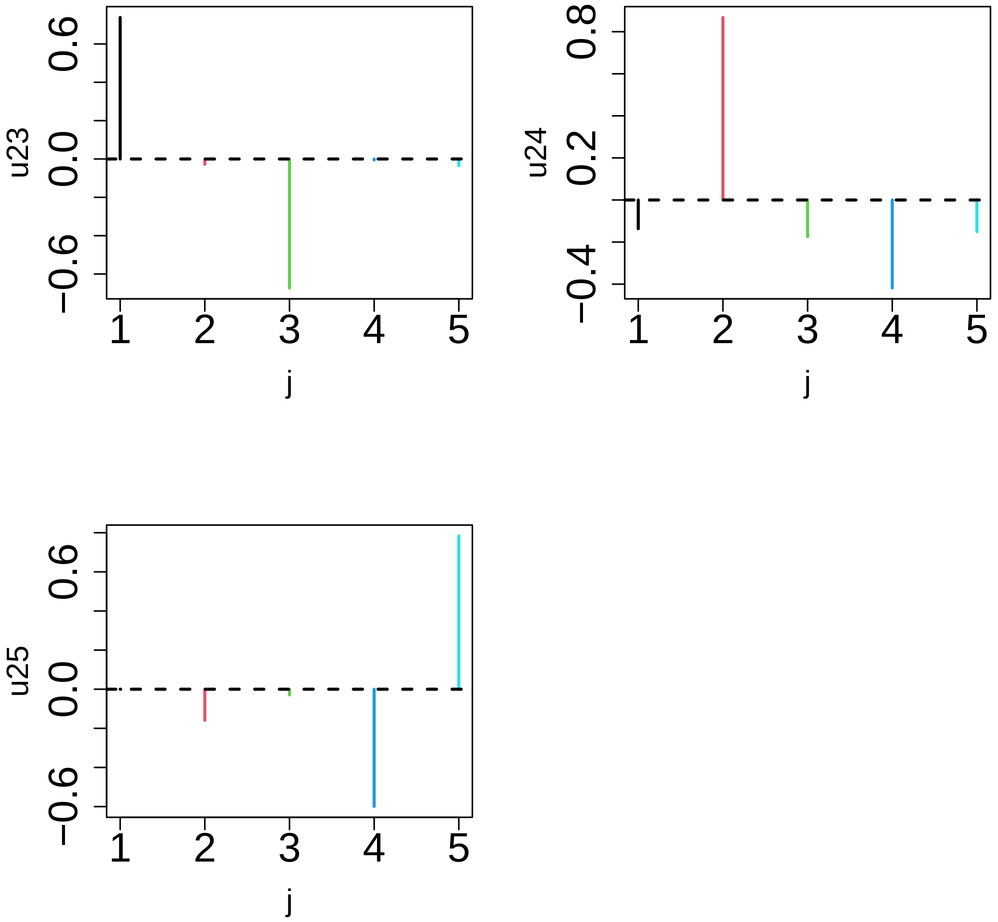


Figure S4: $u_{\ell_2 j}$ not shown in Fig. 1(e)

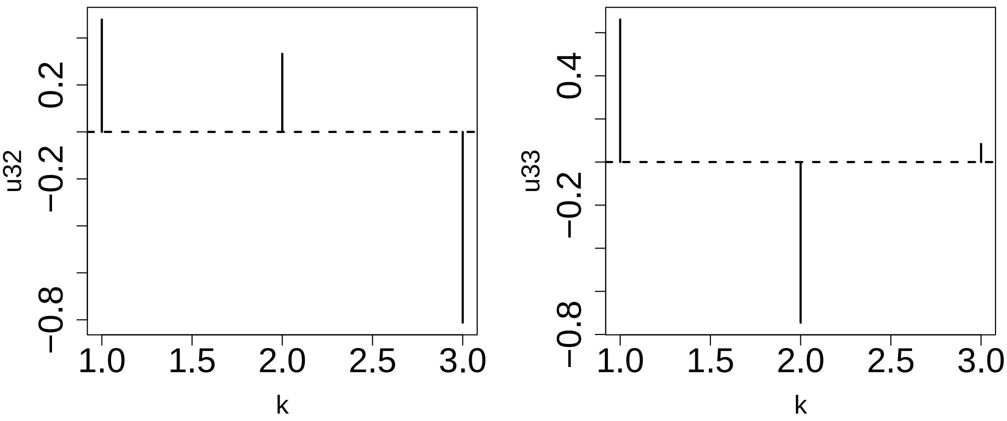


Figure S5: $u_{\ell_3 k}$ not shown in Fig. 1(e)

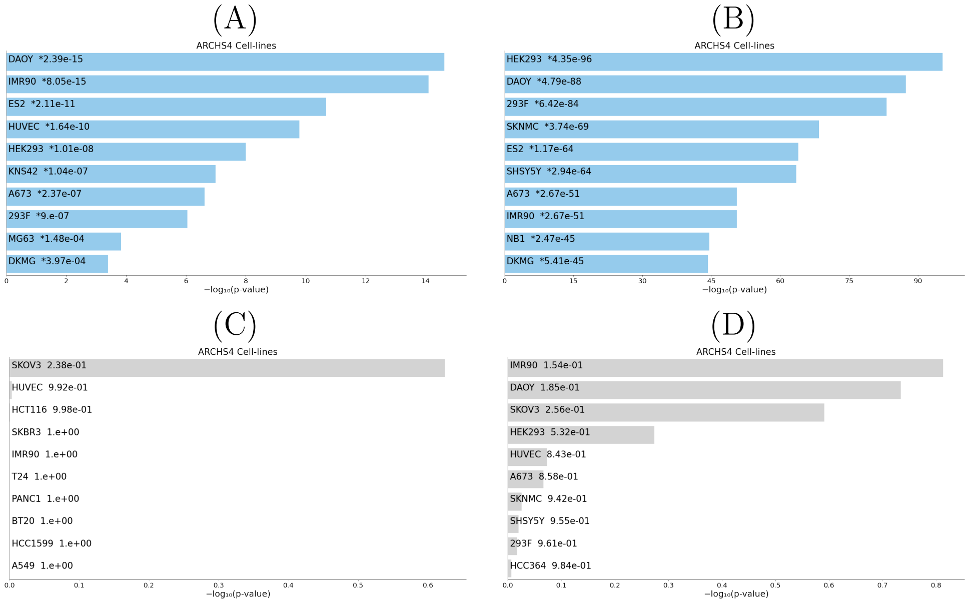


Figure S6: Top ranked 10 cell lines in “ARCHS4 Cell-lines” category of Enricher for genes selected by the proposed method for (A) breast and (B)colon data sets, respectively, and that for those selected by COHCAP for (C) breast and (D) colon data sets, respectively. Colored: raw $P < 0.05$ and asterisked : adjusted $P < 0.05$, respectively.

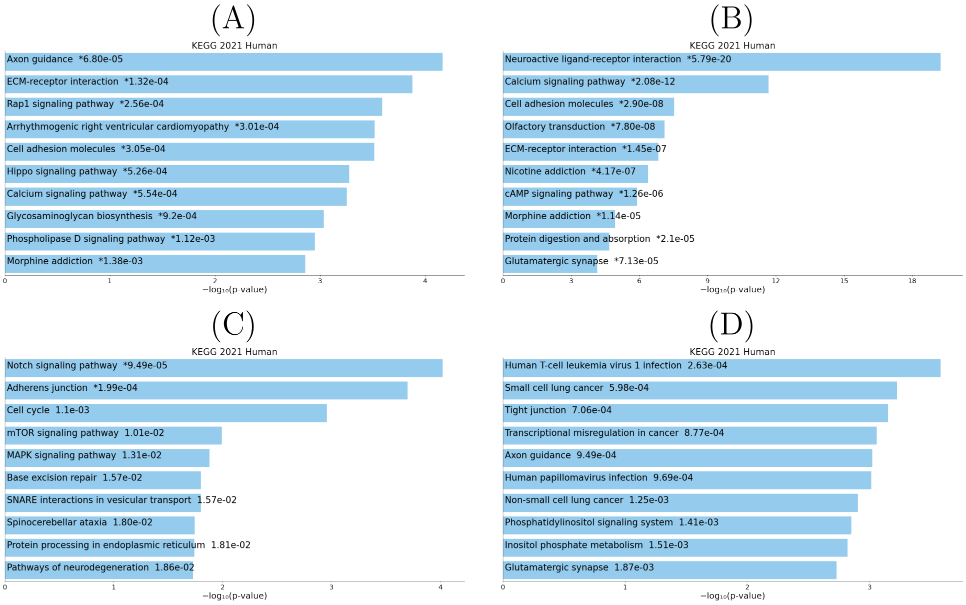


Figure S7: Top ranked 10 pathways in "KEGG 2021 HUMAN" category of Enricher for genes selected by the proposed method for (A) breast and (B)colon data sets, respectively, and that for those selected by COHCAP for (C) breast and (D) colon data sets, respectively. Colored: raw $P < 0.05$ and asterisked : adjusted $P < 0.05$, respectively.

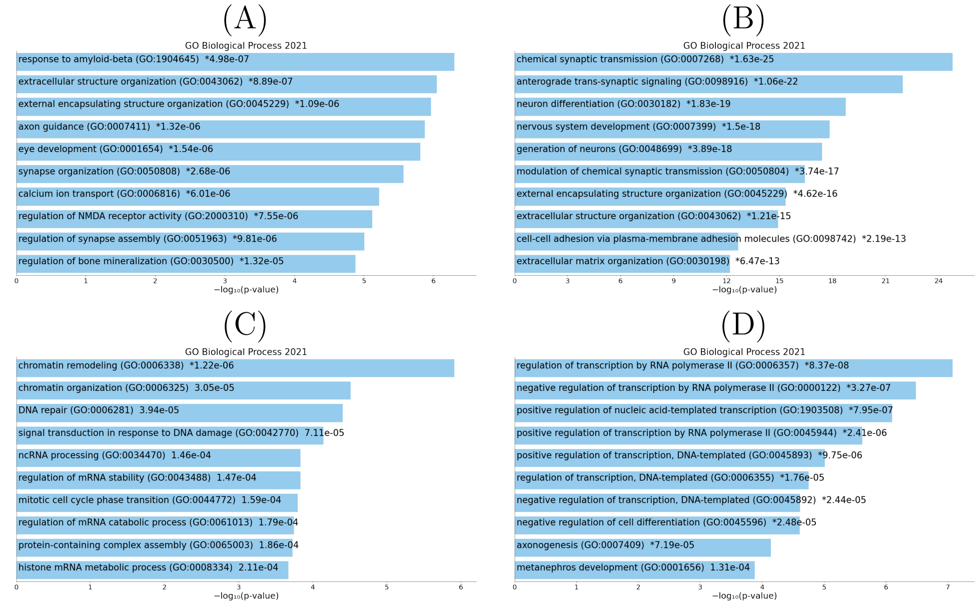


Figure S8: Top ranked 10 biological terms in “GO Biological process 2021” category of Enricher for genes selected by the proposed method for (A) breast and (B)colon data sets, respectively, and that for those selected by COHCAP for (C) breast and (D) colon data sets, respectively. Colored: raw $P < 0.05$ and asterisked : adjusted $P < 0.05$, respectively.

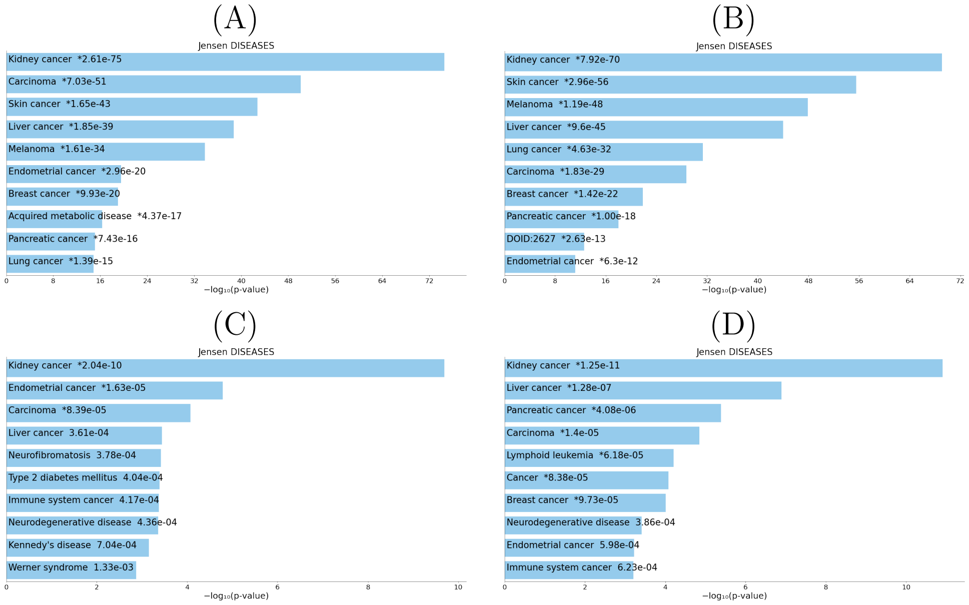


Figure S9: Top ranked 10 diseases in "Jensen Diseases" category of Enricher for genes selected by the proposed method for (A) breast and (B)colon data sets, respectively, and that for those selected by COHCAP for (C) breast and (D) colon data sets, respectively. Colored: raw $P < 0.05$ and asterisked : adjusted $P < 0.05$, respectively.

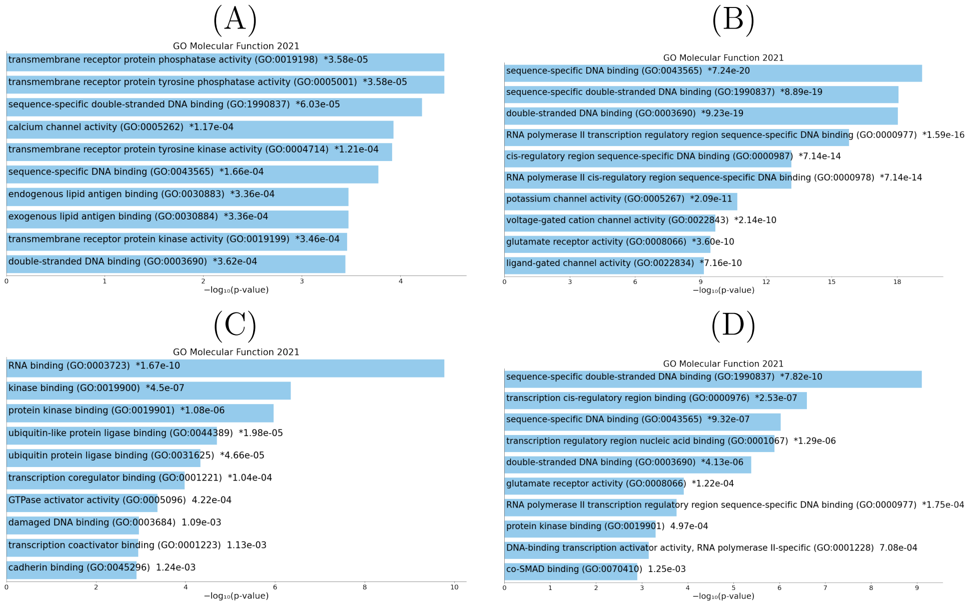


Figure S10: Top ranked 10 functions in "GO Molecular function 2021" category of Enricher for genes selected by the proposed method for (A) breast and (B)colon data sets, respectively, and that for those selected by COHCAP for (C) breast and (D) colon data sets, respectively. Colored: raw $P < 0.05$ and asterisked : adjusted $P < 0.05$, respectively.

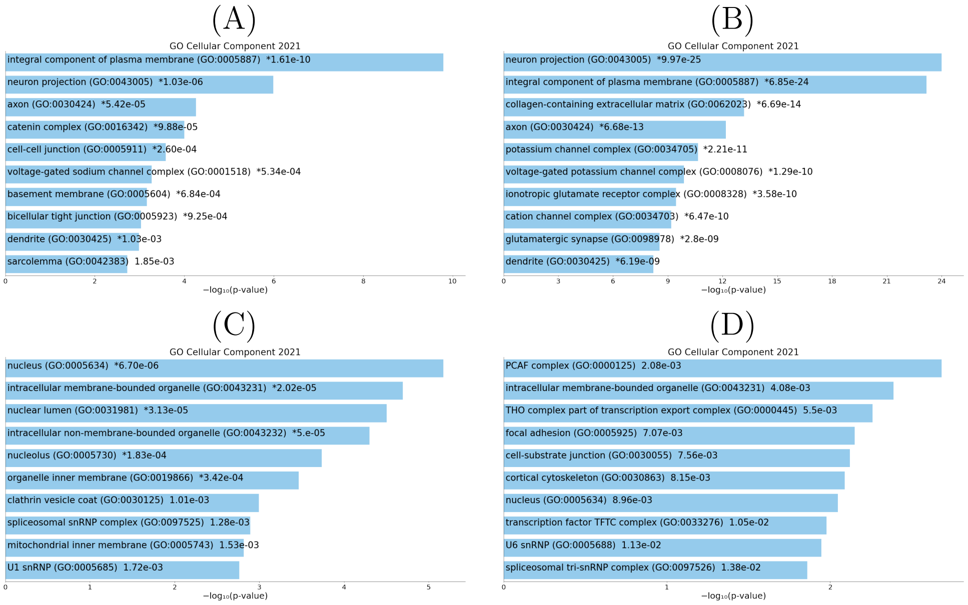


Figure S11: Top ranked 10 components in "GO Cellular Components 2021" category of Enricher for genes selected by the proposed method for (A) breast and (B)colon data sets, respectively, and that for those selected by COHCAP for (C) breast and (D) colon data sets, respectively. Colored: raw $P < 0.05$ and asterisked : adjusted $P < 0.05$, respectively.
