## Supplementary Document(Algorithm) for "Principal component analysis- and tensor decomposition-based unsupervised feature extraction to select more suitable differentially methylated cytosines: Optimization of standard deviation versus state-of-the-art methods"

**Algorithm S1** PCA based unsupervised FE with SD optimization

**Require:** $X \in \mathbb{R}^{N\times M}$:matrix, $L \in \mathbb{R}^{M}$:labels
**Ensure:** $i$:selected feature

**procedure** PCA_BASED_SELECTION$(X, L)$
  $U, V \leftarrow X$ ▷ Compute PC components $U \in \mathbb{R}^{N\times \min(N,M)}, V \in \mathbb{R}^{M\times \min(N,M)}$ from $X$ by PCA
  $P_\ell \leftarrow V, L$ ▷ Compute P-value, $P_\ell$, that assumes the null hypothesis $V[j,\ell]$ is independent of $L_j$
  $P_\ell^{adj} \leftarrow P_\ell$ ▷ Correct $P_\ell$ and get adjusted $P_\ell^{adj}$
  $\ell \leftarrow P_\ell^{adj}$ ▷ Select $\ell$ associated with $P_\ell^{adj} \leq 0.05$
  $\sigma_\ell \leftarrow U[,\ell]$ ▷ Compute standard deviation $\sigma_\ell$ of $U[,\ell]$
  **repeat**
    $P_i \leftarrow U[i,\ell], \sigma_\ell$ ▷ Attribute P-value, $P_i$, to $i$th variable with the null hypothesis that $U[i,\ell]$ obeys Gaussian with $\sigma_\ell$
    $h_n \leftarrow P_i$ ▷ Compute histogram, $h_n$, of $P_i$ at $n$th bin. $(n \leq 100)$
    $P_i^{adj} \leftarrow P_i$ ▷ Correct $P_i$ and get adjusted $P_i^{adj}$
    $\sigma_h \leftarrow h_n$ ▷ Compute standard deviation, $\sigma_h$, of $h_n$ with using only $i$ with $P_i^{adj} > 0.01$
  **until** minimize$\sigma_h$ with regard to $\sigma_\ell$
  $P_i^{adj} \leftarrow P_i$ ▷ Correct $P_i$ and get adjusted $P_i^{adj}$
  $i \leftarrow P_i^{adj}$ **return** $i$ ▷ Select $i$ associated with $P_i^{adj} \leq 0.01$
**end procedure**

**Algorithm S2** TD based unsupervised FE with SD optimization

**Require:** $X \in \mathbb{R}^{N\times M\times K}$:tensor, $L \in \mathbb{R}^{M}, L' \in \mathbb{R}^{K}$:labels
**Ensure:** $i$:selected feature

**procedure** TD_BASED_SELECTION$(X, L, L')$
  $U_1, U_2, U_3, G \leftarrow X$ ▷ Compute core tensor $G \in \mathbb{R}^{N\times M\times K}$ and singular value vetors, $U_1 \in \mathbb{R}^{N\times N}, U_2 \in \mathbb{R}^{M\times M}, U_3 \in \mathbb{R}^{K\times K}$ from $X$ by TD
  $P_\ell \leftarrow U_2, L; P'_\ell \leftarrow U_3, L'$ ▷ Compute P-value, $P_\ell$ and $P'_\ell$, that assumes the null hypothesis $U_2[j,\ell]$ and $U_3[k,\ell]$ are independent of $L_j$ and $L'_k$
  $P_\ell^{adj} \leftarrow P_\ell; P_\ell'^{adj} \leftarrow P'_\ell$ ▷ Correct $P_\ell$ and $P'_\ell$ and get adjusted $P_\ell^{adj}$ and $P_\ell'^{adj}$
  $\ell \leftarrow P_\ell^{adj}, \ell' \leftarrow P_\ell'^{adj}$ ▷ Select $\ell$ and $\ell'$ associated with $P_\ell^{adj}$ and $P_\ell'^{adj} \leq 0.05$
  $\ell'' \leftarrow G[\ell'', \ell, \ell']$ ▷ Select $\ell''$ with the largest absolute value of $G[\ell'', \ell, \ell']$
  $\sigma_{\ell''} \leftarrow U_1[,\ell'']$ ▷ Compute standard deviation $\sigma_{\ell''}$ of $U_1[,\ell'']$
  **repeat**
    $P_i \leftarrow U_1[i,\ell''], \sigma_{\ell''}$ ▷ Attribute P-value, $P_i$, to $i$th variable with the null hypothesis that $U_1[i,\ell'']$ obeys Gaussian with $\sigma_{\ell''}$
    $h_n \leftarrow P_i$ ▷ Compute histogram, $h_n$, of $P_i$ at $n$th bin. $(n \leq 100)$
    $P_i^{adj} \leftarrow P_i$ ▷ Correct $P_i$ and get adjusted $P_i^{adj}$
    $\sigma_h \leftarrow h_n$ ▷ Compute standard deviation, $\sigma_h$, of $h_n$ with using only $i$ with $P_i^{adj} > 0.01$
  **until** minimize $\sigma_h$ with regard to $\sigma_{\ell''}$
  $P_i^{adj} \leftarrow P_i$ ▷ Correct $P_i$ and get adjusted $P_i^{adj}$
  $i \leftarrow P_i^{adj}$ **return** $i$ ▷ Select $i$ associated with $P_i^{adj} \leq 0.01$
**end procedure**
